# Truly tiny acoustic biomolecules for ultrasound imaging and therapy

**DOI:** 10.1101/2023.06.27.546773

**Authors:** Bill Ling, Bilge Gungoren, Yuxing Yao, Przemysław Dutka, Cameron A. B. Smith, Justin Lee, Margaret B. Swift, Mikhail G. Shapiro

## Abstract

Nanotechnology offers significant advantages for medical imaging and therapy, including enhanced contrast and precision targeting. However, integrating these benefits into ultrasonography has been challenging due to the size and stability constraints of conventional bubble-based agents. Here we describe bicones, truly tiny acoustic contrast agents based on gas vesicles, a unique class of air-filled protein nanostructures naturally produced in buoyant microbes. We show that these sub-80 nm particles can be effectively detected both in vitro and in vivo, infiltrate tumors via leaky vasculature, deliver potent mechanical effects through ultrasound-induced inertial cavitation, and are easily engineered for molecular targeting, prolonged circulation time, and payload conjugation.

## INTRODUCTION

Ultrasound is a powerful modality for medical diagnostics and treatment due to its noninvasive nature, real-time imaging capabilities, and widespread accessibility^1^. Incorporation of nanomaterial probes could significantly enhance these benefits, enabling immune evasion, molecular targeting, extravasation, and multifunctional strategies for improved contrast and drug delivery^2,3^. Although such probes have proven successful in modalities such as nuclear and magnetic resonance imaging^4^, designing truly nanoscale agents for ultrasound continues to pose a challenge. Conventional agents, formulated as lipid-shelled gas microbubbles, are typically limited by surface tension to sizes larger than 1 µm^5^, while proposed submicron agents based on bubbles^6^, droplets^7^, phase-change^8,9^, and gas-trapping particles^10–12^ remain relatively large (>200 nm) and can be difficult to prepare.

In this study, we introduce bicones, truly tiny acoustic reporters based on gas vesicles (GVs), a class of air-filled protein nanostructures assembled by certain aquatic microbes for buoyancy regulation^13,14^. GVs comprise a corrugated protein shell of varying thickness (1-3 nm) that excludes liquid water while allowing dynamic gas exchange, creating a stable nanoscale pocket of air^15,16^. Acoustic waves strongly scatter at this air-liquid interface, enabling GVs to serve as contrast agents^14,17–20^ and reporter genes^21–24^ for ultrasound imaging. GVs are highly versatile, with physical and acoustic properties that are easily tuned by functionalizing the protein shell or by modifying their constituent genes^17,25^. Additionally, GVs can serve as therapeutic platforms by stimulating ultrasound mechanotherapy^26^ and photodynamic therapy^27^. However, their largest dimensions typically exceed 200 nm, putting them on the larger end of biological nanomaterials.

GV formation begins as a soluble nucleation complex, progresses into a biconical structure, and ultimately elongates into a cylindrical shape^15,16^. We hypothesized that we could produce ultrasmall particles by inhibiting GV growth at the bicone stage. In this study, we test this idea and establish the fundamental properties and capabilities of GV bicones for potential applications in ultrasound imaging and therapy, evaluating their imaging contrast, tumor uptake capacity, and interactions with focused ultrasound. Additionally, we devise surface engineering strategies to extend circulation time, target specific cells, and carry protein cargo. Using these truly tiny acoustic biomolecules, we aim to enable further integration of ultrasonography with nanomedicine.

## RESULTS

### Bicones are truly tiny ultrasound contrast agents

We hypothesized that we could disrupt GV growth by deleting gvpN, a protein essential for their elongation^28–30^. We modified a previously described bacterial GV cluster containing structural proteins from *Anabaena flos-aquae* and chaperones from *Bacillus megaterium*^22^ and expressed this construct in *E. coli* (**Fig. 1a**). As hypothesized, transmission electron microscopy (TEM) showed uniformly small, biconical particles (**Fig. 1b**). Further deletions of gvpC, gvpR, gvpT, and gvpU did not affect morphology (**Fig. S1**). Cryogenic electron microscopy (cryo-EM) showed that the bicone shell comprised two low-pitch helices beginning at each tip and converging in the center (**Fig. 1c**), consistent with the structure of GVs^16^. Individual particles had an average diameter of 39.7 ± 0.5 nm, length of 72.3 ± 1.1 nm, and enclosed gas volume of 0.023 aL (76.3% of total volume, assuming a 2-nm thick shell^13,31^) (**Fig. 1d, Table S1**). Bicones were colloidally stable in phosphate-buffered saline (PBS), with a hydrodynamic diameter of 58.4 ± 0.5 nm, as measured by dynamic light scattering (DLS) (**Fig. 1e**), and zeta potential of −34.9 ± 1.2 mV (**Fig. 1f**). The volume of a typical bicone is approximately 93 times smaller than the average GV used in most ultrasound experiments and 17 times smaller than a lentiviral capsid (**Fig. 1g, Table S1**).

**Figure 1.**
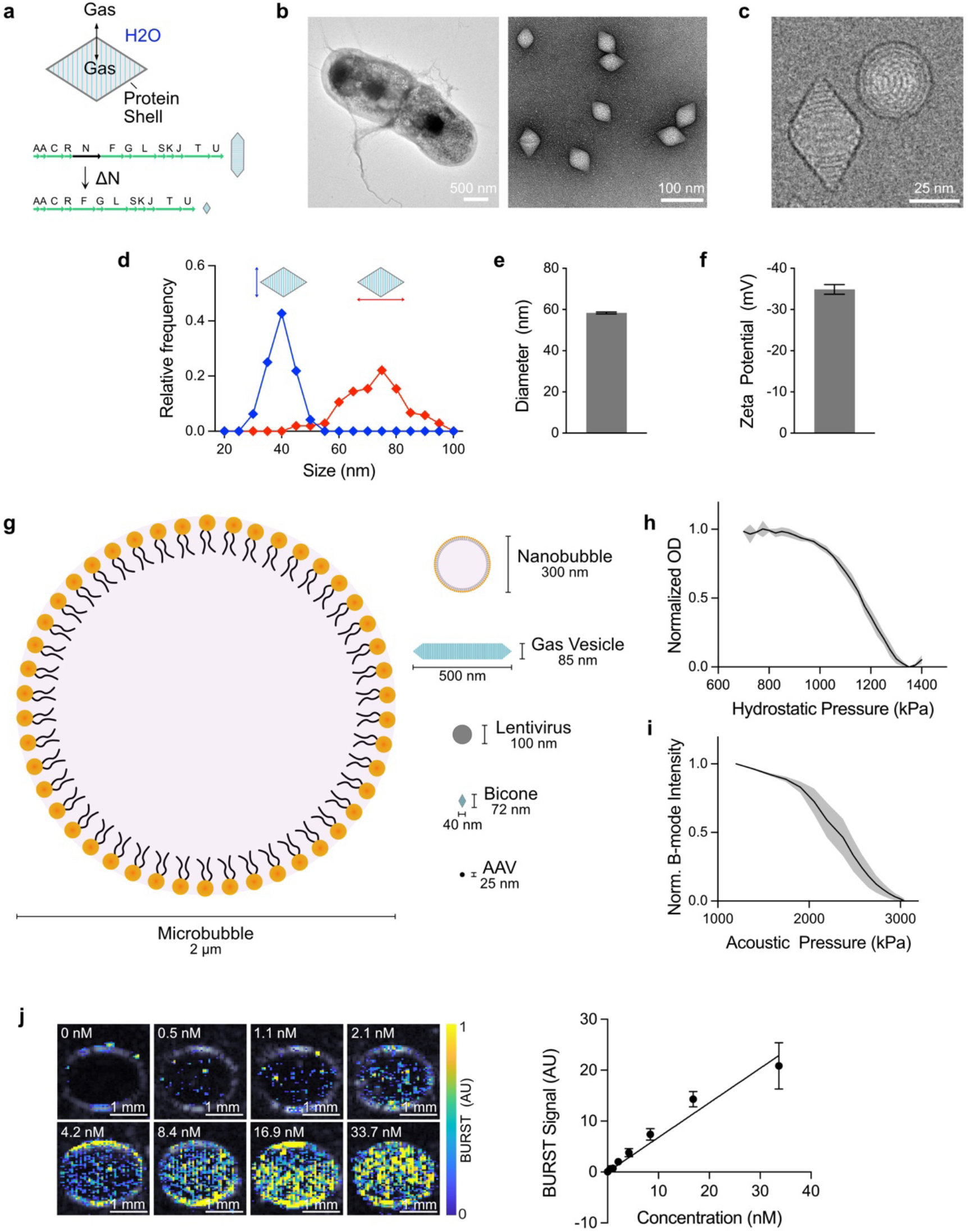
Bicones are truly nanoscale ultrasound contrast agents. **a)** Diagram of bicone structure and gene cluster. Gas can dynamically exchange through the bicone shell, forming a thermodynamically stable pocket of air. Deletion of the gvpN gene prevents GVs from growing beyond the bicone stage. **b)** Representative TEM images of bicones in *E. coli* (left, scale bar, 500 nm) and after purification (right, scale bar, 100 nm). **c)** Representative cryo-EM image of purified bicones showing the helical shell structure. Scale bar, 25 nm. **d)** Distribution of lengths and diameters of individual particles, measured manually from cryo-EM images. *N* = 100. **e)** Intensity-weighted DLS measurements of bicones in PBS. *N* = 35. Error bars, ± SEM. **f)** Zeta potential measurements. *N* = 6. Error bars, ± SEM. **g)** Diagram comparing the dimensions of a typical microbubble, nanobubble, GV, lentivirus, bicone, and adeno-associated virus (AAV). **h)** Pressurized absorbance spectroscopy measurements of bicones in PBS, normalized to starting OD. *N* = 6. Thick line, mean; shaded area, ± SEM. **i)** B-mode intensity of bicones embedded in an agarose phantom following exposure to pulses at the indicated acoustic pressure, normalized to the starting value. *N* = 6. Thick line, mean; shaded area, ± SEM. **j)** Representative BURST images of bicones embedded in an agarose phantom at concentrations between 0 nM and 33.7 nM, overlaid on a B-mode image to show sample outlines. Scale bars, 1 mm. **k)** Quantification of BURST signal within samples from panel J. Data were fit by linear regression, slope = 0.86, r^2^ = 0.5674. *N* = 12. Error bars, ± SEM.

Bicones are expected to be highly resistant to external pressure, as shell robustness is inversely related to diameter^32^. We tested this using pressurized absorbance spectroscopy, which measures optical density under increasing hydrostatic pressure to identify the threshold when GVs collapse and lose their internal gas content and consequent ability to scatter light^33^. Optical density decreased gradually between 0.9 MPa and 1.3 MPa, with a midpoint at 1.16 MPa (**Fig. 1h**). To assess collapse under acoustic pressure, we embedded bicones in an agarose phantom and acquired B-mode ultrasound images at a center frequency of 15 MHz, with increasing transducer driving voltage. Contrast diminished sharply at 2 MPa and was completely erased at 3 MPa, with a midpoint of approximately 2.4 MPa (**Fig. 1i**). These collapse thresholds rank bicones as the sturdiest GV variant we have developed (**Fig. S2**).

Having confirmed that our transducers could collapse bicones, we visualized them using BURST imaging^21^. This method maximizes sensitivity and specificity for GVs by capturing the transient signals generated during acoustic collapse. BURST signal correlated linearly with concentration and was reliably detected at 0.5 nM, the lowest concentration tested (**Fig. 1j-k**).

### Bicones enable visualization of tumors

We next investigated the ability of bicones to produce ultrasound contrast *in vivo*. We hypothesized that bicones could be visualized in mouse xenografts, as sub-100 nm nanoparticles are expected to accumulate in cancerous tissue through leaky vasculature^34–36^. To test this hypothesis, we intravenously (IV) administered 25 pmol bicones into nude mice bearing subcutaneous U-87 MG tumors and performed BURST imaging after 1 h. We chose this dose to maximize delivery by exploiting potential clearance saturation mechanisms^37^. We expected to see contrast within a several-millimeter band of depth due to transducer focusing and the high collapse pressure of bicones. Indeed, BURST signal was detected as a punctate band within the tumor, which was absent during a subsequent acquisition, confirming its specificity to intact particles (**Fig. 2a, S3**). We validated tumor accumulation in a separate group of mice by measuring the fluorescence of tumors resected 2 h after injection of bicones labeled with a near-infrared dye (**Fig. 2b-c**).

**Figure 2.**
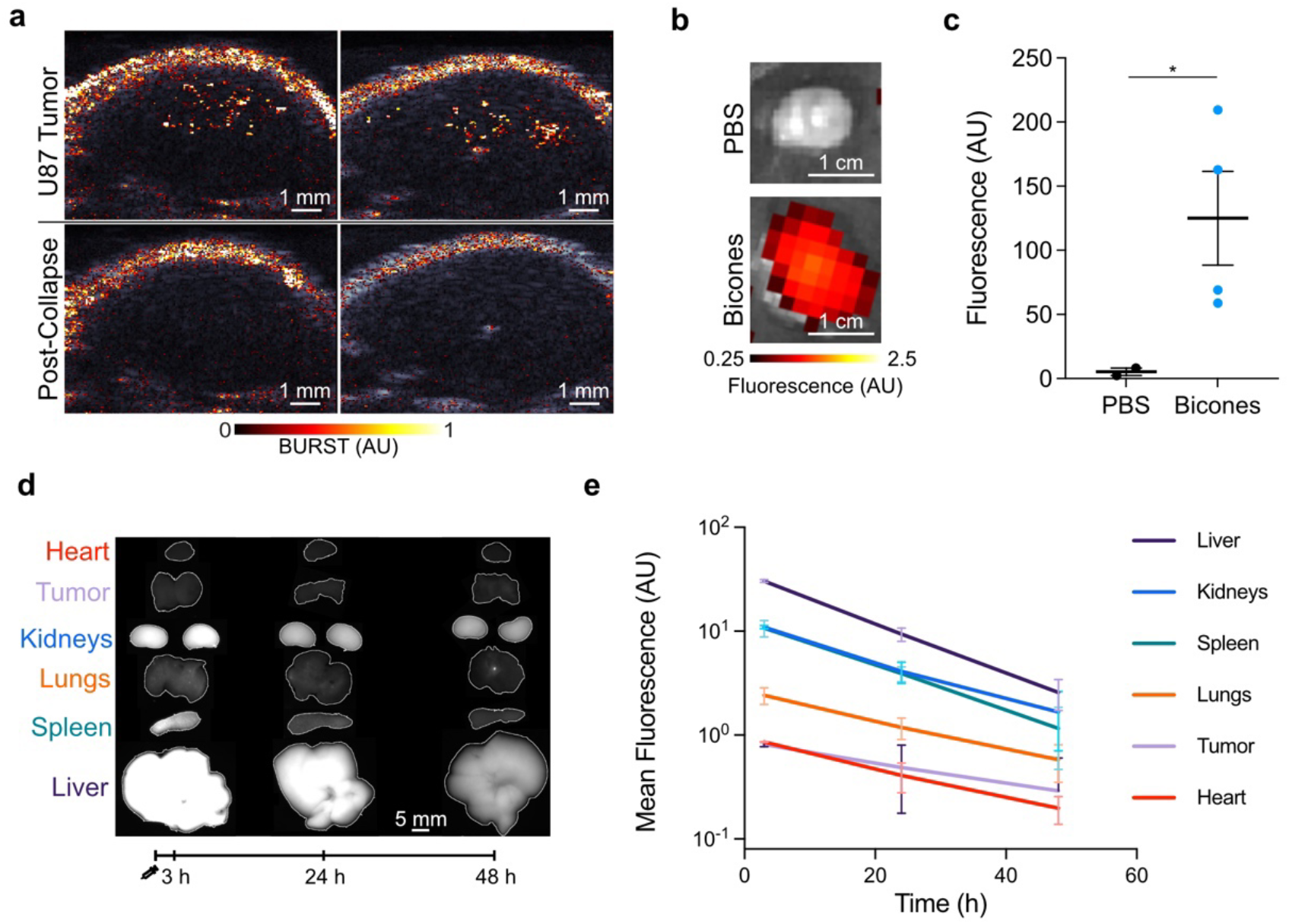
Tumor imaging and biodistribution of bicones. **a)** Representative BURST images of subcutaneous U-87 MG tumors 1 h after intravenous injection of bicones, overlaid on a B-mode image of the anatomy. Bottom row shows a second acquisition at the same location, confirming signal was specific to intact bicones. Images here are from one of three mice (see **Fig. S3**). Scale bars, 1 mm. **b)** Representative fluorescence images of U-87 MG tumors resected 2 h after IV injection of PBS (*N* = 2) or bicones labeled with a near-infrared fluorescent dye (*N* = 4), overlaid on a grayscale photograph to show tissue boundaries. Scale bars, 1 cm. **c)** Fluorescence intensity of tumors in panel B. Errors bars, ± SEM. Welch’s t-test, (*, p < 0.05). **d)** Representative fluorescence images of mouse organs excised at the specified time after IV injection of bicones bearing a far-red fluorescent dye. Organs are outlined in white. Scale bar, 5 mm. **e)** Mean fluorescence intensity within each organ from panel D. *N* = 4 at each time point. Error bars, ± SEM. Data for spleen and tumor are partially obscured by kidneys and heart, respectively.

To examine biodistribution, we acquired fluorescence images of mouse organs excised at predetermined intervals following IV injection of bicones labeled with a farred fluorescent dye (**Fig. 2d-e**). Fluorescence was highest in the liver, spleen, and kidneys. At 3 h post-injection, mean intensities in these organs were 30.5, 10.7, and 11.0, respectively, decreasing to 9.4, 3.9, and 4.1 after 24 h and further to 2.6, 1.2, and 1.7 after 48 h. This gradual reduction in fluorescence over time indicates active elimination of bicones from the body, with the high kidney signal suggesting lysosomal degradation followed by renal excretion^18,38^. Fecal excretion may also contribute to elimination, as shown by biliary and intestinal accumulation of GVs from *H. salinarum*^39^, though we did not examine these organs in this study. Compared to larger GVs^18,39^, bicones showed considerably lower uptake in the spleen and lungs, consistent with the ability of smaller particles to bypass filtration by these organs^40^. Overall, our data demonstrate that bicones are remarkably small acoustic biomolecules that can be easily produced in bacteria and detected both *in vitro* and *in vivo*.

### Bicones can seed inertial cavitation

Having demonstrated bicone accumulation in tumors, we next explored their potential for therapeutic applications. Specifically, we investigated their capacity to seed inertial cavitation under focused ultrasound (FUS)^26^. This phenomenon can be used to precisely disrupt tissue, promote drug penetration, and eliminate diseased cells^41^. Inertial cavitation involves the growth of a bubble—often nucleated by air carried in synthetic agents^11^ or released during GV collapse^26^—through coalescence across several acoustic cycles, culminating in a high-energy implosion (**Fig. 3a**). Bubbles undergoing this process generate distinct broadband acoustic emissions^26^.

**Figure 3.**
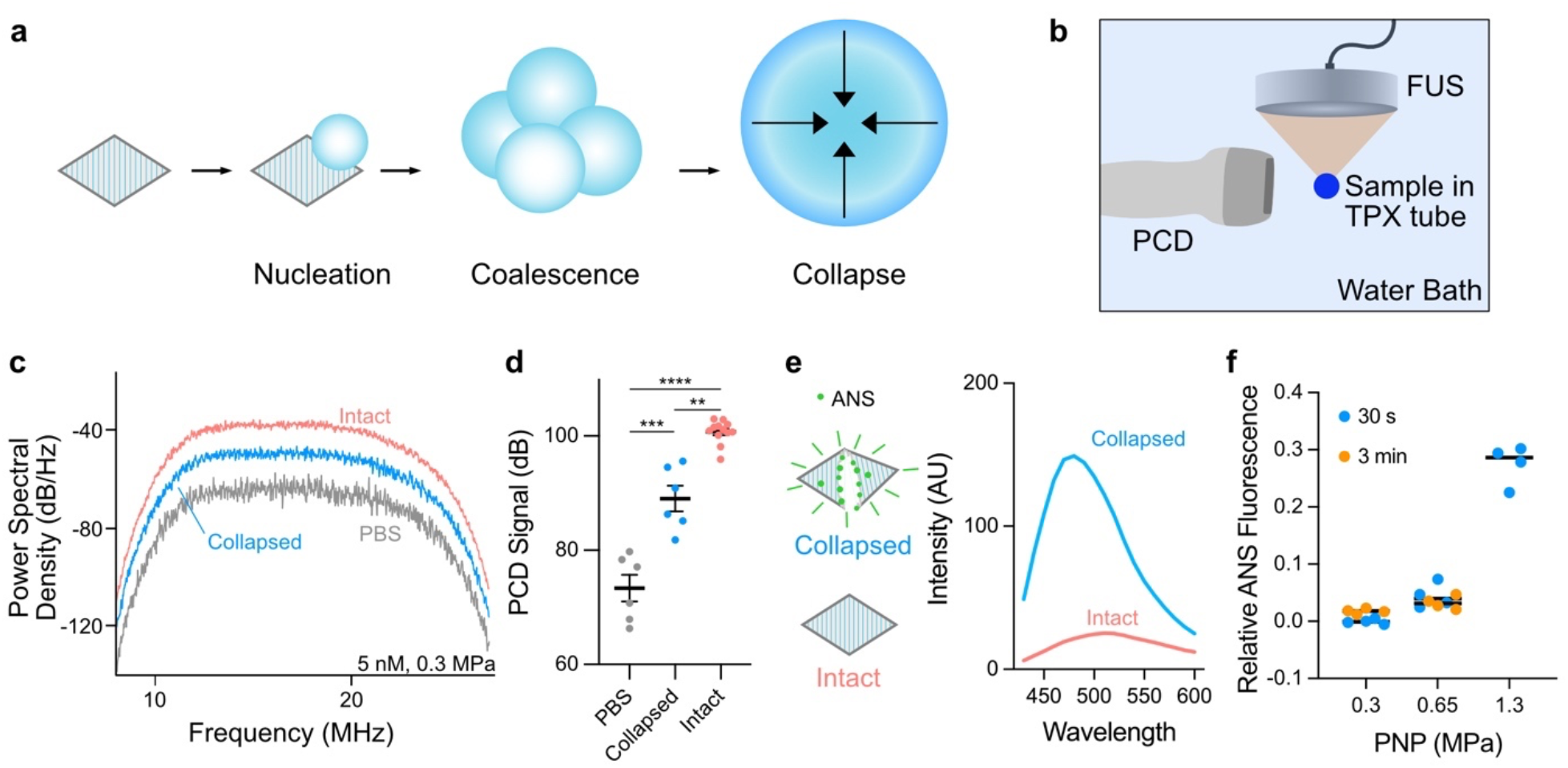
Bicones can seed inertial cavitation. **a)** Mechanism of bicone-mediated inertial cavitation. Bubbles are nucleated by the gas content of bicones and grow through coalescence across multiple acoustic cycles, culminating in a high-energy implosion. **b)** Diagram of experimental setup to measure acoustic emissions induced by FUS. Samples contained in acoustic impedance-matched TPX tubes were insonated with a 330-kHz FUS transducer and responses were detected by an orthogonally positioned imaging transducer. **c)** Representative power spectral density of emissions from PBS, pre-collapsed, and intact bicones (5 nM) insonated with a single 30 cycle FUS pulse at 0.3 MPa PNP. **d)** Mean PCD signal of samples from panel C, calculated by integrating spectra from 8 MHz to 27 MHz. PBS, *N* = 6; collapsed, *N* = 6; intact, *N* = 12. Error bars, ± SEM. Welch’s t-test (**, p < 0.01; ***, p < 0.001; ****, p < 0.0001). **e)** Left: ANS dye fluoresces upon binding the exposed hydrophobic interior of the bicone shell. Right: Representative emission spectra (ex. 380 nm) of intact and collapsed bicone suspensions containing 100 µM ANS. **f)** Relative ANS fluorescence at 480 nm of bicones after exposure to 30 s or 3 min of FUS pulses at various PNP, normalized to fluorescence of intact and pre-collapsed samples. *N* = 4. Error bars not shown.

To test the ability of bicones to seed inertial cavitation, we insonated samples with a 330 kHz FUS transducer and recorded acoustic emissions using an orthogonally positioned imaging transducer as a passive cavitation detector (PCD) (**Fig. 3b**). We applied FUS to bicone suspensions in acoustic impedance-matched polymethylpentene (TPX) tubes, chosen to minimize attenuation. At a peak negative pressure (PNP) of 0.3 MPa, intact bicones (5 nM) seeded broadband emissions of greater power than buffer (PBS) control (**Fig. 3c-d**). This pressure and frequency correspond to a mechanical index (MI) of 0.52 MPa MHz^-1/2^, well below the accepted safety limit for diagnostic ultrasound^42^. Collapsed bicones were partially effective in seeding cavitation, presumably by stabilizing bubbles with their exposed hydrophobic surfaces. Similar responses were observed at a PNP of 0.5 MPa (MI = 0.87 MPa MHz^-1/2^, **Fig. S4**). Cavitation seeded by bicones was comparable in power to that of *Anabaena* GVs at the same gas fraction (**Fig. S5**). Insonation with 670 kHz FUS at 0.3 MPa PNP yielded emission spectra with enhanced power at harmonic multiples of the transmitted frequency, suggesting the presence of stably oscillating bubbles^26^ (**Fig. S6**).

Given the low pressures at which we observed cavitation—below the bicones’ hydrostatic and acoustic collapse thresholds—we wondered whether the particles remain intact while seeding cavitating bubbles. To independently track bicone collapse, we used the solvatochromic dye 8-anilinonaphthalene-1-sulfonic acid (ANS)^43,44^, which we reasoned would fluoresce more brightly upon binding to the hydrophobic interior of the bicone shell exposed during the collapse process. As expected, bicones that were collapsed in the presence of 100 µM ANS either by bulk bath sonication (at 40 kHz) or hydrostatic pressure greatly increased the dye’s fluorescence (**Fig. 3e, S7**). Fluorescence intensity at 480 nm correlated linearly with the concentration of collapsed bicones (**Fig. S8**).

Using ANS as a readout, we quantified collapse in 4.2 nM bicone suspensions after extended 330 kHz FUS insonation with 100 µs pulses at a 500 Hz pulse repetition frequency. We calculated the percentage of collapsed particles based on fluorescence intensity relative to intact and bath-sonicated samples. Less than 5% of bicones collapsed after either 30 s or 3 min of sonication at 0.3 MPa or 0.65 MPa PNP (**Fig. 3f**). In comparison, 43% of *Anabaena* GVs collapsed after just 20 ms of pulses at 0.65 MPa (**Fig. S9**). At the higher pressure of 1.3 MPa, 3 min of sonication was sufficient to collapse over 25% of the bicones (**Fig. 3f**). Although 1.3 MPa is below the acoustic collapse threshold of bicones (**Fig. 1i**), applied pressure is expected to be amplified in the near vicinity of cavitating bubbles. Taken together, these data suggest that bicones can seed cavitation at low applied pressure and may be able to do so repeatedly by remaining intact.

### Polymer coated bicones have enhanced circulation

Having identified potential diagnostic and therapeutic applications for bicones, we next developed strategies to optimize their longevity in the bloodstream. Specifically, we sought to coat bicones with methoxypolyethylene glycol (mPEG), a polymer widely used for immune evasion^45,46^. We focused on azide-alkyne cycloadditions^47–49^, as these bio-orthogonal reactions are compatible with GVs^50^ and enable convenient access to any azide-functionalized substrate. We attached alkyne groups to lysines on the bicone surface and conjugated 10 kDa azide-mPEG using a copper-catalyzed alkyne-azide cycloaddition (CuAAC) reaction^51^ (**Fig. 4a**). Consistent with the addition of a PEG layer, hydrodynamic diameter increased from 58.4 ± 3.0 nm to 97.5 ± 12.9 nm (**Fig. 4b**) and zeta potential neutralized from −34.9 ± 1.2 mV to −5.2 ± 1.6 mV (**Fig. 4c**).

**Figure 4.**
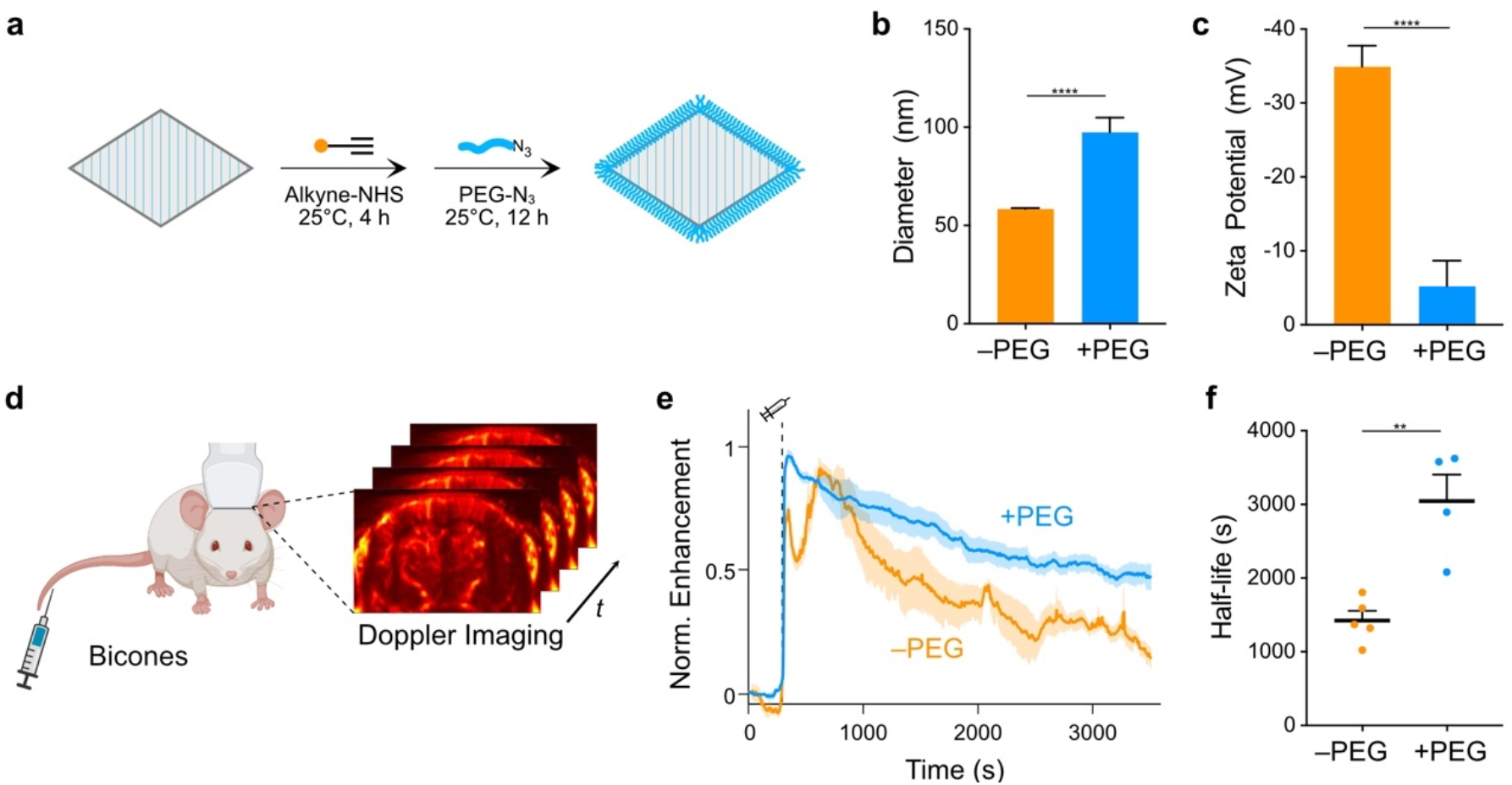
Extension of bicone circulation time. **a)** Diagram of PEGylation protocol. Lysines on the bicone surface were functionalized with alkyne groups and attached to 10 kDa mPEGs through a CuAAC reaction. **b-c)** DLS (**b**) and zeta potential (**c**) measurements of PEG-coated bicones. Data from Fig. 1 shown for comparison (–PEG). *N* = 5. Error bars, ± SEM. Welch’s t-test (****, p < 0.0001). **d)** Diagram of experimental setup to measure circulation time. Intravascular bicones were visualized by ultrafast power Doppler ultrasound imaging of the brain. **e)** Normalized time courses of hemodynamic contrast enhancement following IV injection of bicones. Time of injection (300 s) is indicated by a dashed line. Thick lines, mean; shaded area, ± SEM. *N* = 4-5. **f)** Circulation half-life calculated by fitting time courses in panel E to an exponential decay function. Error bars, ± SEM. Welch’s t-test, (**, p < 0.01).

We tested the effectiveness of this modification by measuring apparent circulation time in BALB/c mice. We visualized circulating bicones with ultrafast power Doppler ultrasound imaging, leveraging their ability to enhance blood flow contrast^18,19^. Targeting a single coronal plane in the brain, we acquired images at a center frequency of 15.625 MHz and frame rate of 0.25 Hz (**Fig. 4d**). After a 5 min baseline, we IV-injected 13 pmol bicones and monitored the ensuing changes in hemodynamic signal (**Fig. 4e**). Unmodified bicones produced signal enhancement time courses with two distinct peaks, while PEGylated bicones only generated a single peak before returning to baseline monotonically. Fitting to an exponential decay function showed that unmodified bicones circulated with a half-life of 1424 ± 132 s, while PEGylated bicones circulated with a half-life of 3045 ± 362 s (**Fig. 4f**). Notably, this is an order of magnitude longer than most synthetic agents which typically circulate for only several minutes^1^.

### Biochemical functionalization for targeting

Molecular targeting is another key capability for molecular imaging agents. To demonstrate the potential for such targeting with bicones, we functionalized their surfaces with cyclic iRGD peptides, which bind to integrins that are overexpressed in certain pathological tissues^52,53^. We coupled dibenzocyclooctyne (DBCO) and a far-red fluorophore to amines on the bicone shell at molar ratios of 1000 and 2500 per particle, respectively. We then attached azide-iRGD through a strain-promoted alkyne-azide cycloaddition (SPAAC) reaction^48,50^ (**Fig. 5a**). We incubated the human cancer cell lines U-87 MG and HT-1080 with these particles for 2 h at 37°C. After rinsing the cells with PBS to remove unbound bicones, we observed binding by confocal microscopy (**Fig. 5b**). We further validated these results by analyzing the cells via flow cytometry (**Fig. 5c, S10**). On average, 34.7% of U-87 MG and 60.3% of HT-1080 cells exposed to iRGD-bicones exhibited significant increases in fluorescence, compared to less than 3% of cells exposed to untargeted bicones (**Fig. 5d**). Notably, binding was enhanced using a maximum stoichiometry of just 10 peptides per particle and increased with ligand loading (**Fig. S11**).

**Figure 5.**
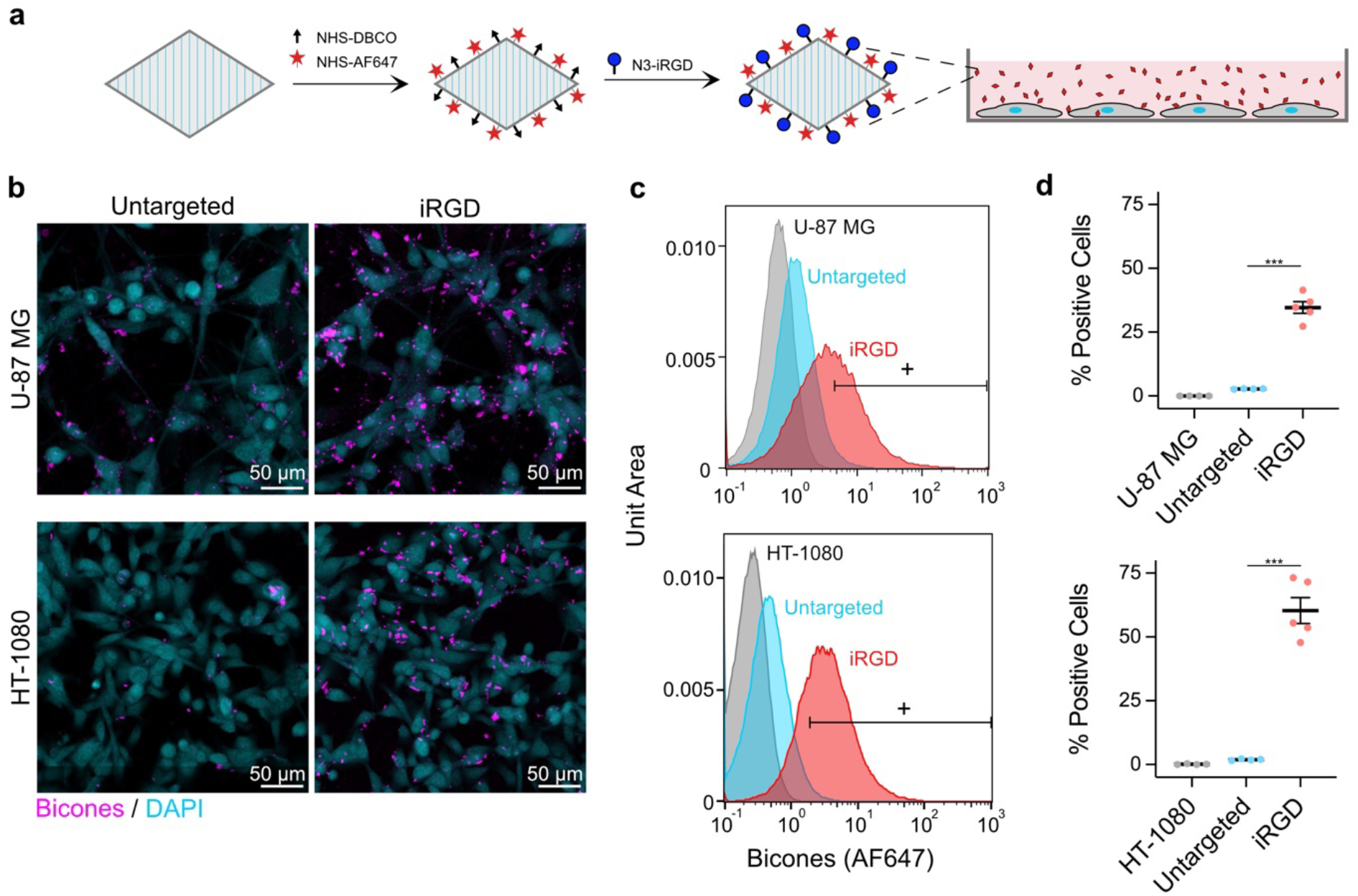
Molecular targeting with bicones. **a)** Diagram of iRGD conjugation protocol. Bicones were dually-labeled with DBCO and a fluorescent dye, then functionalized with cyclic iRGD peptides by a SPAAC reaction. Purified particles were incubated *in vitro* with human cancer cells. **b)** Representative confocal microscopy images of cells after incubation with bicones. Scale bars, 50 µm. **c)** Representative flow cytometry histograms of U-87 MG (top) and HT-1080 (bottom) cells incubated with untargeted or iRGD-conjugated bicones. Bicone-positive cells were gated based on fluorescence in samples incubated with PBS only. **d)** Percentage of bicone-positive cells in histograms from panel D. *N* = 4-5. Error bars, ± SEM. Welch’s t-test, (***, p < 0.001).

### Genetic functionalization

As genetically encoded agents, bicones can also be functionalized by modification of their constituent proteins, providing a simple and scalable approach that is orthogonal to chemical bioconjugation. Though previously achieved in full GVs via peptide fusion to GvpC^17,25,54^, SDS-PAGE analysis showed that bicones lack this protein (**Fig. S12**). To identify alternative options, we appended a FLAG tag^55^ to each gene in the GV cluster and tested for association of the fusion protein with purified particles by anti-FLAG dot blot (**Fig. 6a**). GvpA and GvpG did not tolerate fusions (no bicones were produced), while staining from the GvpC and GvpF fusions was weak. However, GvpJ and GvpS produced strong staining. These proteins share homology with the primary structural protein GvpA, suggesting that they may be incorporated directly into the GV shell. Indeed, washing the particles with 6 M urea to remove surface-bound proteins^25^ did not affect GvpJ or GvpS staining (**Fig. S13**).

**Figure 6.**
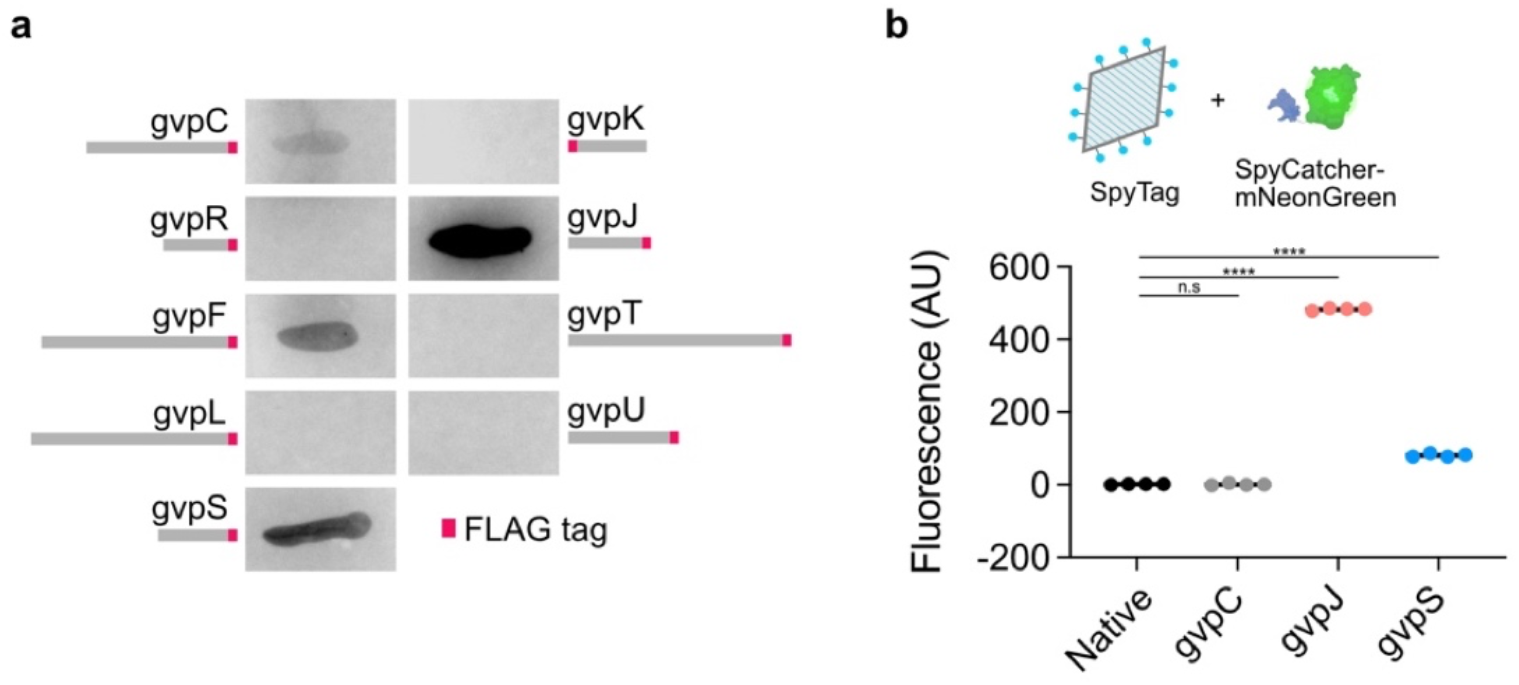
Genetic functionalization of bicones. **a)** Representative images of dot blots. A FLAG tag (shown in red) was appended to the end of each GV gene and incorporation of the fusion protein into purified bicones was assessed using an anti-FLAG antibody. **b)** Top: Purified bicones, unmodified or with SpyTag appended to gvpC, gvpJ, or gvpS, were reacted with the fluorescent protein, SpyCatcher-mNeonGreen. Bottom: Mean fluorescence intensity of purified particles after conjugation. *N* = 4. Error bars, ± SEM. Welch’s t-test, (****, p < 0.0001; n.s, p ≥ 0.05).

Using these new genetic handles, we developed a modular system to covalently conjugate recombinant proteins to the bicone surface. We fused SpyTag, a 13-residue peptide that spontaneously forms a covalent bond with its partner SpyCatcher domain under physiological conditions^56^, to the ends of GvpC, GvpJ, and GvpS. We reacted the purified particles with the model payload SpyCatcher-mNeonGreen^25^ (SC-mNG), chosen for easy quantitation of binding through fluorescence (**Fig. 6b**). After removing free protein, neither the GvpC mutant nor unmodified bicones exhibited any increase in fluorescence. In contrast, fluorescence from the GvpJ and GvpS fusions increased significantly, with the largest increase for GvpJ. When SC-mNG was reacted with similar constructs expressed on full-size GVs with GvpC^22^, all three fusion proteins exhibited strong fluorescence (**Fig. S14**). Our results indicate that GvpJ and GvpS are molecular handles for bicone functionalization which can be used in conjunction with the SpyTag-SpyCatcher system to facilitate convenient and modular bioconjugation; these handles were also effective in full-size GVs. Together with the chemical approaches described above, bicones have versatile competence for molecular functionalization.

## DISCUSSION

Our findings establish bicones as versatile, truly nanoscale agents for ultrasound imaging and cavitation. Bicones help bridge the gap between ultrasound contrast agents and nanomedicine, potentially unlocking applications beyond the capabilities of large, unstable bubble-based agents. Easily produced in bacteria, their small size allows for extended blood circulation and passive tumor uptake through leaky vasculature. Bicones also serve as nuclei for inertial cavitation and notably do so at low pressures without collapsing. With these properties, bicones could enable enhanced imaging and mechanotherapy in tumors and mucosal^57^, pulmonary^58^, and lymphatic systems^59^, where larger particles are rapidly sequestered.

The functionality of bicones is further enhanced by the ease with which their surfaces can be modified. This confers additional capabilities, including immune evasion, molecular targeting, and payload conjugation. Furthermore, their compatibility with CuAAC and SPAAC chemistry provides a convenient pathway to tailor ligands to individual applications.

However, two key improvements are necessary to enhance the utility of bicones. First, they should be made compatible with non-destructive imaging methods. While we used BURST imaging for its sensitivity, visualizing bicones without causing collapse would streamline diagnostic applications. This issue could be addressed by engineering shell proteins to allow bicones to buckle reversibly under acoustic pressure and generate nonlinear echoes that can be captured using specialized pulse sequences^25,60^. Second, a comprehensive investigation of cavitation processes is required. While our data suggest that bicones can seed cavitation without collapse, the extent of their capacity for repeated bubble nucleation in various media and *in vivo* requires more comprehensive analysis. As cavitation seeds, bicones could offer simpler production and functionalization processes compared with synthetic agents^11,61^.

Over the long term, bicone production and purification processes should be optimized. Lengthy centrifugation steps are impractical at larger scales, and alternatives such as tangential flow filtration or chromatography could be explored. Formal studies of toxicity and biocompatibility should also be conducted in tandem to inform efforts to refine purification processes. While the doses we used did not cause adverse health effects in mice, residual bacterial proteins and antibody responses could negatively impact long-term performance.

In conclusion, bicones could allow approaches previously exclusive to nuclear and magnetic resonance imaging to be translated into the cost-effective and accessible realm of ultrasonography, highlighting the immense potential of these truly tiny particles.

## METHODS

### Chemicals

All chemicals were purchased from Sigma Aldrich and used without further purification unless otherwise noted.

### Bicone preparation

Plasmid construction: The original bicone construct (A2C-∆ N) was prepared by deleting gvpN from the full GV gene cluster (available on Addgene as plasmid #106473) via KLD mutagenesis using enzymes from New England Biolabs and primers from IDT. Briefly, this plasmid contained gvpAAC from *Anabaena flos-aquae* and gvpRFSKJTU from *Bacillus megaterium* downstream of a T7 promoter on a pET28a backbone. Variants were similarly created. FLAG and SpyTag variants included a short linker (GGSG) before the peptide tag. Primer sequences are included in **Table S2**.

Expression and purification: Plasmids were transformed into *E. coli* strain BL21(DE3) (New England Biolabs) and grown on plates overnight at 37 °C (LB agar, 2%w/v glucose, 100 µg mL^-1^ kanamycin). Starter cultures were prepared by inoculating a large number of colonies into 4 mL Miller’s LB supplemented with 50 µg mL^-1^ kanamycin and 2%w/v glucose and grown at 37°C for 3 h. Large-scale cultures were then prepared by 1:100 dilution of the starter culture in 50 mL LB containing kanamycin and 0.2%w/v glucose. Expression was induced after 2.5 h (OD600 ∼0.6) by addition of 10 µM isopropyl-β-D -1-thiogalactopyranoside (IPTG, Teknova), grown overnight at 37 °C, and harvested by centrifugation in 50-mL conical tubes at 450xg (12 h, 4 °C). Excess LB was removed by vacuum filtration through binder-free glass microfiber filters (Whatman GF/F grade, Cytiva). Cells were lysed by incubation in 4 mL SoluLyse (Genlantis) supplemented with 100 µg DNaseI (Roche) for 5 h at room temperature. Lysates were then combined and clarified by centrifugation at 600xg for 2 h at 4 °C. Bicones were enriched from the supernatant and exchanged into PBS by 5 rounds of overnight centrifugation at 800xg, 4 °C followed by replacement of the subnatant with fresh PBS. Concentrations were measured with a Bio-Rad Quick Start Bradford protein assay based on bovine serum albumin standards.

### Particle characterization

Dynamic light scattering: A disposable semi-micro polystyrene cuvette (VWR) containing 300 µL of a 2 µg mL^-1^ bicone suspension was placed in a Brookhaven Instruments ZetaPALS particle analyzer, and particle size was determined with the ZetaPALS Particle Sizing software using an angle of 90°, thin shell setting, run length of 15 s, and 6 runs per sample. Measurements were recorded as mean intensity-weighted diameter.

Zeta potential: An electrode (SZP, Brookhaven Instruments) was inserted into a mixture of 50 µL of 200 µg mL^-1^ bicones in PBS and 1.5 mL Milli-Q water in a disposable plastic cuvette (SCP, Brookhaven Instruments). Measurements were performed with the ZetaPALS Zeta Potential software using the Smoluchowski model, based on 5 runs of 15 cycles.

Pressurized absorbance spectroscopy: Hydrostatic collapse pressure measurements were performed as previously described^33^. Briefly, 350 µL of a 200 µg mL^-1^ bicone suspension in PBS was loaded into a flow-through quartz cuvette (Hellma Analytics). Hydrostatic pressure was applied from a nitrogen gas source through a single valve pressure controller (Alicat Scientific). Pressure was ramped from 700 kPa to 1400 kPa in 25 kPa increments with a 7 s equilibration period prior to measurement of absorbance at 600 nm with a spectrophotometer (OceanOptics STS-VIS). Bicones collapsed at 1.5 MPa were used as a blank.

TEM: Samples in PBS were diluted 1:10 in water and spotted onto Formvar/carbon 200 mesh copper grids (Ted Pella) that were glow discharge treated (Emitek K100X). The grids were negatively stained with 1% uranyl acetate. Images were acquired with a Thermo Fisher FEI Tecnai 120 keV T12 LaB6 electron microscope equipped with a Gatan Ultrascan 2k x 2k CCD camera and Gatan Digital Micrograph data collection software. Images were processed with Fiji^62^.

CryoEM: Images were acquired as previously described^63^. Gold grids (C-Flat 2/2 − 2Au) were glow discharge treated (Pelco EasiGlow). Freshly purified bicone samples (0.5 mg mL^-1^) were spotted onto the grids and flash-frozen using a Mark IV Vitrobot (Thermo Fisher) (4 °C, 100% humidity, blot force 3, blot time 6 s). Images were acquired on a 300 kV Titan Krios microscope (Thermo Fisher) equipped with an energy filter (Gatan) and a K3 6k x 3k direct electron detector (Gatan) using SerialEM software^64^. Images were loaded into Fiji for manual measurement of particle dimensions.

### Bicone functionalization

Stoichiometries were calculated using 800 gvpA per particle and 1.65e11 particles per µg protein (**Table S1**). These estimates were based on a conical geometric model with average dimensions as measured by cryo-EM, assuming a protein density of 1.4 mg/µL and a 2-nm thick shell composed purely of gvpA.

PEGylation: Alkyne-bicones were prepared by mixing purified bicones with propargyl-N-hydroxysuccinimide ester (prepared as a 250 mM stock solution in DMSO) at a 120:1 molar ratio of ester to gvpA and gently rotated at room temperature for 4 h. The particles were then purified by 4 rounds of overnight centrifugation at 800xg, 4 °C. The CuAAC reaction was prepared by combining 22 mg m-PEG-azide (10 kDa, BroadPharm), 44 µL DMSO, 4.8 µL PBS, 21.5 µL aminoguanidine hydrochloride (prepared as 11.11 mg mL^-1^ PBS), and 44 µL of a 1:1 mixture of BTTAA (Click Chemistry Tools, 77.4 mg mL^-1^ PBS) and copper sulfate pentahydrate (7.4 mg mL^-1^ water). Once the mPEG was fully dissolved, 1.5 mL of a 400 µg mL^-1^ alkyne-bicone suspension and 25 µL sodium ascorbate (59.4 mg mL^-1^ in PBS, freshly made within 1 h of reaction) were added, and the mixture was rotated slowly overnight at room temperature. PEG-bicones were purified by 4 rounds of centrifugation.

iRGD conjugation: Azide-iRGD peptide was custom synthesized by Thermo Fisher (sequence: azide-PEG4-GGSGGS[C]RGDKGPD[C]). DBCO-bicones were prepared by reacting purified bicones with Alexa Fluor 647-succinimidyl ester (Invitrogen, prepared as 10 mM stock in DMSO) and dibenzocyclooctyne-sulfo-N-hydroxysuccinimide ester (DBCO-Sulfo-NHS ester, Click Chemistry Tools, prepared as 10 mM stock in DMSO) at a 2500:1:1000 molar ratio of dye:particle:DBCO. The reaction was allowed to proceed for 4 h at room temperature before purification by 2 rounds of centrifugation. Azide-iRGD peptide (prepared as 1 mg mL^-1^ stock in PBS) was added at a 2:1 molar ratio of peptide to particle, incubated overnight at 4 °C with gentle rocking, then purified by 3 rounds of centrifugation.

### In vitro ultrasound imaging

Images were acquired using a 128-element linear array probe (L22-14vX, Verasonics) with a center frequency of 18 MHz and elevation focus of 8 mm. The transducer was connected to a programmable ultrasound scanner (Verasonics Vantage 128) operating on Vantage 4.4.0 software.

Phantoms were cast from 1% agarose in PBS using custom printed molds containing pairs of 2-mm diameter cylindrical wells. Samples were gently mixed 1:1 with 2% low-melt agarose in PBS and quickly loaded into the wells. Phantoms were placed on acoustic absorber material and immersed in PBS to couple the sample with the imaging transducer.

Acoustic collapse measurement: Images were acquired using a conventional B-mode sequence operating at 15.625 MHz with a 40-element aperture. The transducer was positioned such that the wells were at a depth of 8 mm. An automated voltage ramp script was programmed to insonate samples for 10 s at a specified transmit voltage before recording a single frame at 1.6 V. Transmit voltage was ramped from 2 V to 20 V in 0.5 V increments. The fraction of intact GVs at each pressure step was calculated based on mean B-mode intensity within each well, with samples collapsed at 25 V as a blank.

BURST imaging: BURST images were acquired using a previously described imaging sequence^21^. Briefly, a conventional B-mode sequence optimized for frame rate was used to acquire 10 baseline frames at a voltage of 1.6 V and 20 collapse frames at 20 V. BURST signal in each well was calculated by integrating mean signal intensity across all collapse frames using the final frame as a blank. For display, the first two collapse frames were processed with a 1-pixel radius median filter. The second frame was then subtracted from the first, and voxels with intensities above a threshold were overlaid on a B-mode image.

### Focused ultrasound-induced cavitation

Passive cavitation detection: Measurements were performed as previously described^26^. Under B-mode guidance, an L22-14vX imaging transducer was positioned 10 mm from the center of a 3D-printed holder inside of a water tank. A focused ultrasound transducer (Sonic Concepts HT-115) mounted on a computer-controlled 3D-translatable stage (Velmex) was positioned orthogonal to the imaging probe and aligned to the center of the holder based on feedback from a needle hydrophone (HNR-1000, ONDA Corporation). To enable passive cavitation detection, the Verasonics system was programmed with a zero-amplitude transmit and synchronized with the FUS pulse.

Bicone suspensions were dispensed into 1.5 mL polymethylpentene (TPX) tubes (Diagenode) and allowed to equilibrate overnight at ambient conditions. For PCD measurements, samples were insonated with a 30-cycle burst of 330 kHz sine waves, and emissions were sampled at 62.5 MHz. Power spectral densities were computed for each individual channel using Welch’s overlapped segment estimation method and averaged across all 128 transducer elements. PCD signal was calculated by integrating each spectrum from 8 MHz to 27 MHz using trapezoidal sums.

Fluorescence detection of collapse: Suspensions comprising 25 µg mL^-1^ bicones and 100 µM 8-anilino-1-napthalenesulfonic acid (ANS) were prepared in TPX tubes. Samples were insonated for the specified amount of time with 30-cycle 330 kHz sine wave bursts repeating at 0.5 kHz (2 ms pulse length, ∼5% duty cycle). Samples were then distributed into 96-well microplates and analyzed with a Tecan Spark microplate reader using an excitation wavelength of 380 nm and scanning the emission spectrum from 420 nm to 600 nm in increments of 10 nm. Fraction of intact bicones was calculated based on fluorescence at 480 nm and normalized relative to samples collapsed in an ultrasonic bath (Branson).

### Genetic functionalization

Dot blot of shell proteins: FLAG tags^55^ (DYKDDDDK) were individually appended to each member of the A2C-∆ N gene cluster and purified as described above. PVDF membranes (Bio-Rad) were wetted in methanol, rinsed in 1x TBS-T (137 mM sodium chloride, 20 mM Tris, 0.1% Tween-20) (diluted from 10x solution from Cell Signaling Technology), and placed on top of a stack of 2 sheets of 3MM Chr chromatography paper (Whatman) soaked in TBS-T and 2 sheets of dry 3MM Chr paper. 5 µg of each bicone sample was quickly spotted onto the membrane and allowed to dry completely. The blot was blocked with 5% blotting-grade blocker (Bio-Rad) in TBS-T for 1 h at room temperature and stained overnight at 4 °C with a rabbit anti-FLAG antibody (1:1000 dilution, Millipore F7425). After removing unbound antibodies by three 5-minute washes with TBS-T, the blot was stained with Goat anti-Rabbit IgG (H+L) Secondary Antibody, DyLight 650 (Invitrogen, diluted 1:2000 in 5% blocker) for 2 h at room temperature. Blots were washed 4 times with TBS-T and visualized with a Bio-Rad ChemiDoc MP system using automated exposure settings.

SpyCatcher conjugation: Bicones containing SpyTag peptides^56^ fused to gvpC, gvpJ, or gvpS were prepared as described above. SpyCatcher-mNeonGreen (SC-mNG) was prepared as previously described^25^. Briefly, SC-mNG was expressed in BL21(DE3), purified by non-denaturing Ni-NTA affinity chromatography (Qiagen), and buffer exchanged into PBS by overnight dialysis through regenerated cellulose tubing (6-8 kDa MWCO, Repligen). To test conjugation, 4 µg of SC-mNG and 40 µg bicones were mixed in a total volume of 150 µL PBS and incubated in the dark for 2 h at room temperature. Samples were purified by 2 rounds of centrifugation prior to measuring fluorescence intensity with a Tecan Spark microplate reader (ex. 485 nm, em. 525 nm).

### Cell culture

U-87 MG (HTB-14) and HT-1080 (CCL-121) cells were ordered from the American Type Culture Collection (ATCC) and cultured on tissue culture-treated flasks in Dulbecco’s Modified Eagle’s Medium (Corning) supplemented with 10% fetal bovine serum (Gibco) and 1% penicillin/streptomycin (Gibco) in a humidified incubator at 37°C and 5% CO_2_.

### Cell-specific targeting with iRGD

Confocal microscopy: Tissue culture-treated 24-well plates fitted with #1.5H coverslips (ibidi) were seeded with 50,000 U87 or HT1080 cells per well. After 24 h, the media was exchanged with 500 µL Fluorobrite DMEM (Gibco) supplemented with 10 mM HEPES and 10 µg iRGD-bicones. Following a 2 h incubation, the cells were gently washed twice with 500 µL Fluorobrite DMEM, then fixed by sequential incubation in 2% formalin (10% neutral buffered formalin diluted 1:5 in Fluorobrite) for 2 min and undiluted formalin for 20 min. Samples were stained with 300 nM DAPI (Thermo Fisher) for 5 min and stored in Fluorobrite. Imaging was performed with a Zeiss LSM800 confocal laser scanning microscope through the 20x objective using parameters that prioritized signal specificity over speed. Data were loaded into FIJI and summed over each z-stack.

Flow cytometry: Tissue culture-treated 6-well plates were seeded with 300,000 U87 or HT1080 cells per well. Once the cells reached ∼80% confluency, culture media was removed, and the wells were gently washed with 1.5 mL PBS. The cells were then incubated with 25 µg iRGD-bicones in 1 mL Fluorobrite DMEM (Gibco) for 2 h at 37°C. Afterwards, the cells were gently washed twice with 1 mL room temperature PBS, detached with 500 µL 0.25% trypsin-EDTA (Gibco), and diluted with 500 µL PBS prior to analysis with a Miltenyi MACSQuant Analyzer 10 flow cytometer. Bicone binding was quantified using the R1 channel. Data processing was performed in FlowJo. Gating strategy is shown in **Fig. S10**.

### In vivo ultrasound imaging

All *in vivo* experiments were performed under protocols approved by the Institutional Animal Care and Use Committee at the California Institute of Technology.

Doppler imaging: Female BALB/cJ mice (8-12 wk, Jackson Laboratory) were mounted on a temperature-controlled platform (Stoelting Co., held at 38.5 °C) in a stereotaxic instrument (Kopf Instruments) under isoflurane anesthesia (1.5%, 1L min^-1^), depilated over the skull using Nair, and coupled to an L22-14vX transducer with a column of ultrasound gel (Aquasonic). The transducer was positioned to capture an entire coronal section at an arbitrary plane, and a short catheter (30G needle connected to PE10 tubing) was inserted into the lateral tail vein. Ultrafast power Doppler images were acquired at 15.625 MHz with a frame rate of 0.25 Hz for 1 h using a previously described pulse sequence^18^. Five minutes after the start of acquisition, 200 µL bicones (400 µg mL^-1^) were manually injected into the tail vein.

Data were loaded into MATLAB and hemodynamic signal was isolated by singular value decomposition^65^ (cutoff of 20). Pixel-wise signal enhancement was calculated as the ratio of intensity at each time point relative to mean intensity over the first 75 frames. Time courses were then extracted by averaging signal enhancement within a manually drawn region of interest encompassing the cortex and smoothed with a 10-unit median filter. To calculate half-life, an exponential decay function was fitted to each time course from its maximum to the end of acquisition.

Tumor accumulation: Subcutaneous flank tumors were implanted in 3-5 week old male Nu/J mice (The Jackson Laboratory) using 3e6 U-87 MG cells suspended in 25% Matrigel matrix (Corning) diluted with DMEM. Once the tumors were at least 5 mm in diameter, 200 µL of a 0.5 mg mL^-1^ bicone suspension was injected through the lateral tail vein. After 1 h, the mice were anesthetized under isoflurane and placed on a temperature-controlled platform. An L22-14vX transducer was coupled to the tumor with a column of ultrasound gel and adjusted such that the center of the tumor was at a depth of 8 mm. BURST images were then acquired as described above.

### Biodistribution

Bicones were fluorescently labeled by incubation with 2500x molar equivalents of Alexa Fluor 647 NHS ester (Invitrogen) at room temperature for 4 h and purified by three rounds of centrifugation. Nude mice bearing subcutaneous U-87 MG tumors were IV injected with 200 µL of labeled bicones and transcardially perfused with 50 mL PBS after a predetermined interval. The heart, lungs, kidneys, liver, spleen, and tumor were extracted and imaged with a Bio-Rad ChemiDoc MP using red epi-illumination and a 695/55 nm filter with an exposure time of 0.2 s. Integrated densities were calculated using the built-in “Analyze Particles” function in FIJI. Tumor accumulation was similarly analyzed using the near-infrared dye indocyanine green NHS ester (AdipoGen). Here, only the tumors were resected, and images were acquired with an IVIS Lumina In Vivo imaging system (PerkinElmer) and analyzed using Living Image software (PerkinElmer).

### Statistical analysis

Sample sizes were chosen based on preliminary experiments to yield sufficient power for the proposed comparisons.

Statistical methods are described in applicable figure captions.

## ACKNOWLEDGMENTS

The authors thank Dr. Andres Collazo and the Caltech Biological Imaging Facility of the Beckman Institute for assistance with confocal microscopy; Dr. Songye Chen and the Caltech Cryo-EM facility for assistance with cryo-EM and TEM; Dr. Lena Gamboa and Port Therapeutics for assistance with IVIS imaging; Dr. Di Wu, Dr. Avinoam Bar-Zion, Dr. Constantine Sideris, and Dr. George Lu for helpful discussions. Parts of Fig. 6 were created with BioRender. This research was supported by the National Institutes of Health (grant R01-EB018975 to M.G.S.). M.G.S. is a Howard Hughes Medical Institute Investigator.

## AUTHOR CONTRIBUTIONS

B.L. and M.G.S. conceptualized the research. B.L. and B.G. prepared and characterized all bicone variants. P.D. conducted cryo-EM experiments. B.L., Y.Y, and C.A.B.S. conducted the cavitation experiments. B.L. and J.L conducted flow cytometry experiments. B.L. conducted *in vivo* experiments with assistance from M.B.S. B.L. and M.G.S. wrote the manuscript with input from all other authors.

M.G.S. supervised the research.

## DATA AND MATERIALS AVAILABILITY

All gas vesicles, plasmids, data, and code are available from the authors upon reasonable request.

