## Supplementary Information for "Truly tiny acoustic biomolecules for ultrasound imaging and therapy"

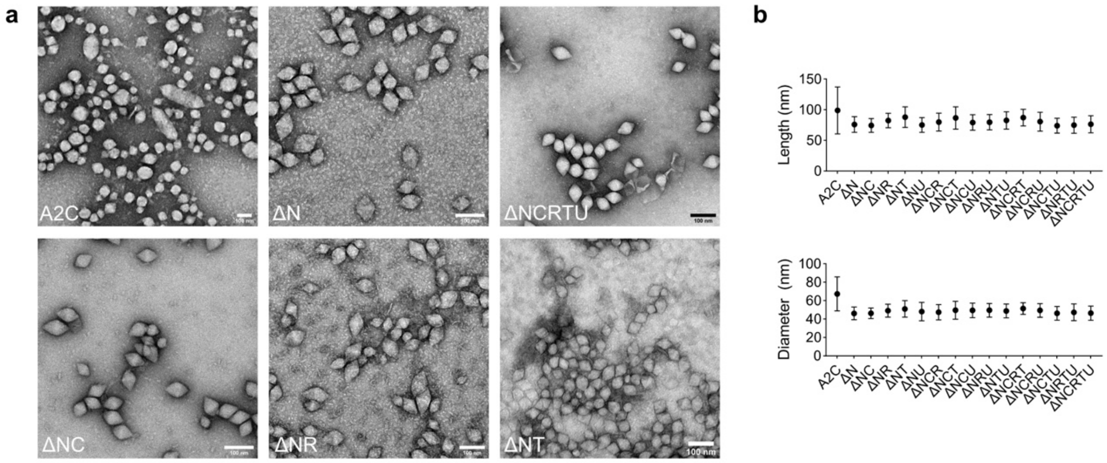


**Figure S1**. Impact of gene deletions on GV morphology. **a)** Representative TEM images of GVs expressed using the indicated gene cluster. A2C represents the full GV cluster[1]. Deletions are indicated by Δ. Scale bars, 100 nm. **b)** Mean length and diameter of individual particles measured in images from panel A. *N* = 50-100 particles. Error bars, ± SD.

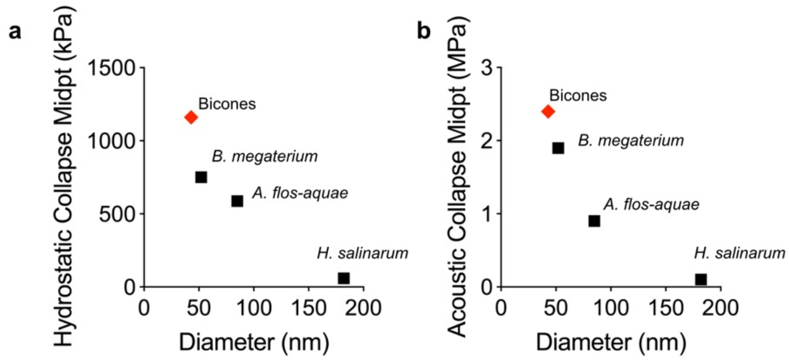


**Figure S2.** Comparison of hydrostatic (**a**) and acoustic (**b**) collapse midpoints for bicones relative to GVs isolated from *H. salinarum* and *A. flos-aquae*[2], or recombinantly expressed in *E. coli* using a cluster native to *B. megaterium*[3].

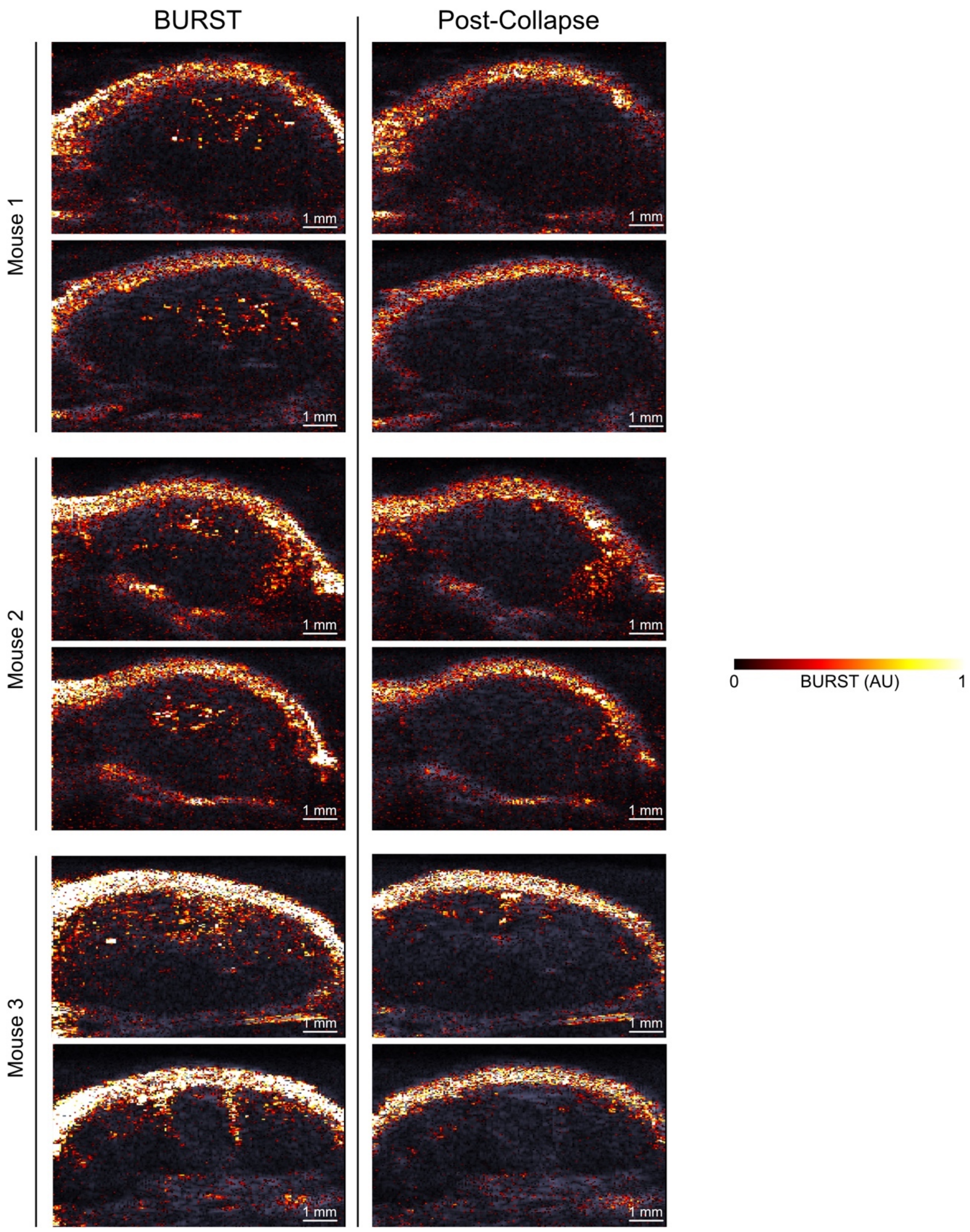


**Figure S3.** BURST images of U-87 MG tumors acquired 1 h after IV injection of bicones, overlaid on a B-mode image to show anatomy. A second acquisition (Post-Collapse) at the same location was used to verify that signal was specific to intact bicones. Two sets of images were acquired at different planes within each tumor. *N* = 3. Scale bars, 1 mm.

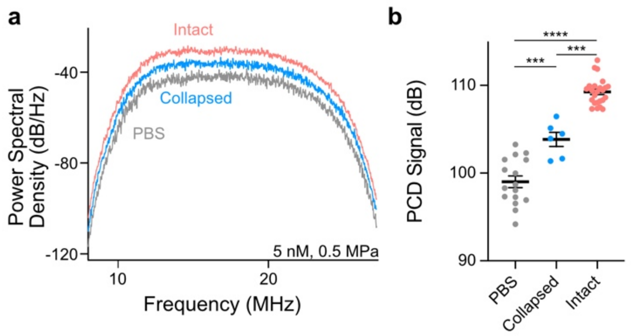


**Figure S4. a)** Representative power spectral density of emissions from PBS, pre-collapsed, and intact bicones (5 nM) insonated with a single 30 cycle FUS pulse at 0.5 MPa PNP. **b)** Mean PCD signal of samples from panel A, calculated by integrating spectra from 8 MHz to 27 MHz. PBS, $N = 16$; collapsed, $N = 6$; intact, $N = 24$. Error bars, ± SEM. Welch's t-test (***, $p < 0.001$; ****, $p < 0.0001$).

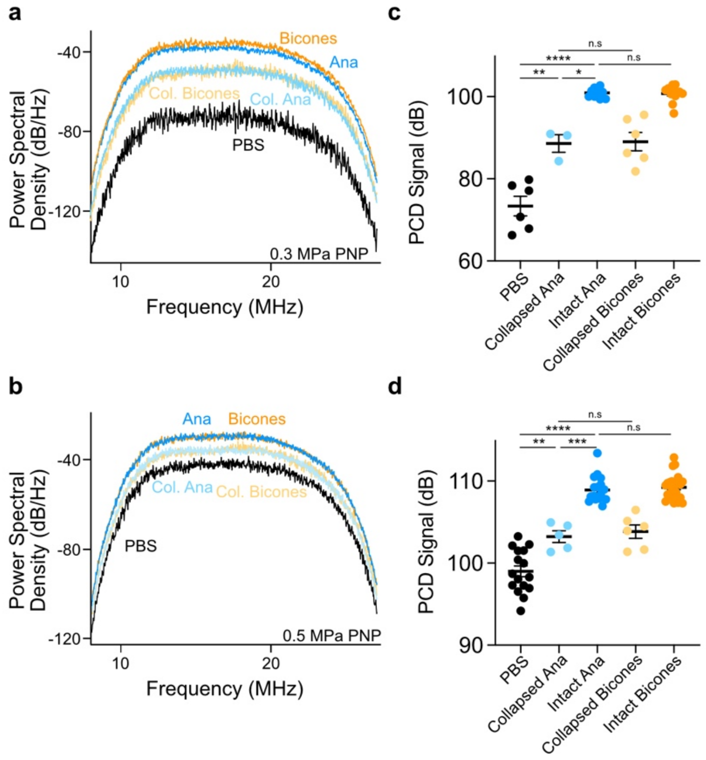


**Figure S5.** Bicones produce comparable PCD signal to *Anabaena* GVs at the same gas volume. **a-b)** Representative power spectral densities of acoustic emissions following insonation of bicones (4.2 nM) and *Anabaena* GVs (OD 0.25) with a single FUS pulse at 0.3 MPa PNP (**a**) or 0.5 MPa PNP (**b**). **c-d)** Mean PCD signal following insonation at 0.3 MPa PNP (**c**) or 0.5 MPa PNP (**d**). Spectra were integrated from 8-27 MHz. $N = 3\text{-}24$. Error bars, ± SEM. Welch's t-test, (*, $p < 0.05$; **, $p < 0.01$; ***, $p < 0.001$; ****, $p < 0.0001$; n.s, $p \geq 0.05$).

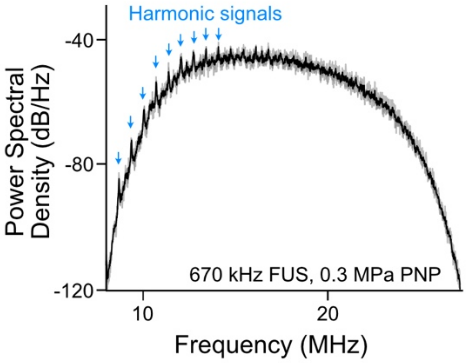


**Figure S6.** Power spectral density of emissions from intact bicones (5 nM) insonated with a single 60 cycle 670 kHz FUS pulse at 0.3 MPa PNP. *N* = 3. Individual trials are shown in gray, and the mean is shown in black. Harmonic signals indicative of stable cavitation are annotated with blue arrows.

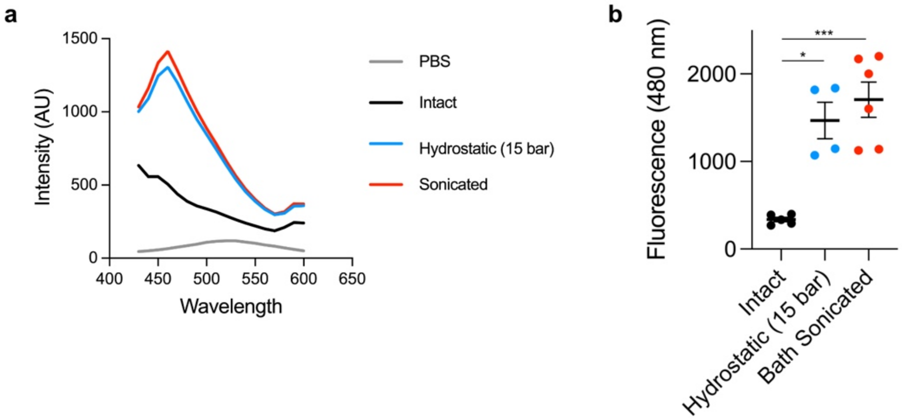


**Figure S7.** ANS fluoresces in the presence of collapsed bicones. **a)** Representative fluorescence emission spectra (ex. 380 nm) of bicone samples with 100 μM ANS following collapse by bath sonication or hydrostatic pressure. **b)** Fluorescence intensity at 480 nm in spectra from panel A. *N* = 4–6. Error bars, ± SEM. Welch's t-test (*, $p < 0.05$; ***, $p < 0.001$).

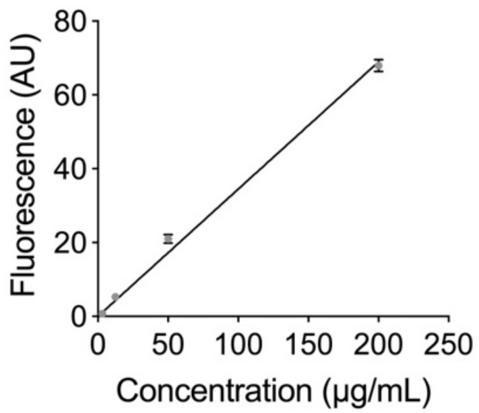


**Figure S8.** Fluorescence of suspensions containing 100 µM ANS mixed with the indicated concentration of collapsed bicones. Data were fit by linear regression, slope = 0.3447, $r^2$ = 0.99. *N* = 4. Error bars, ± SEM.

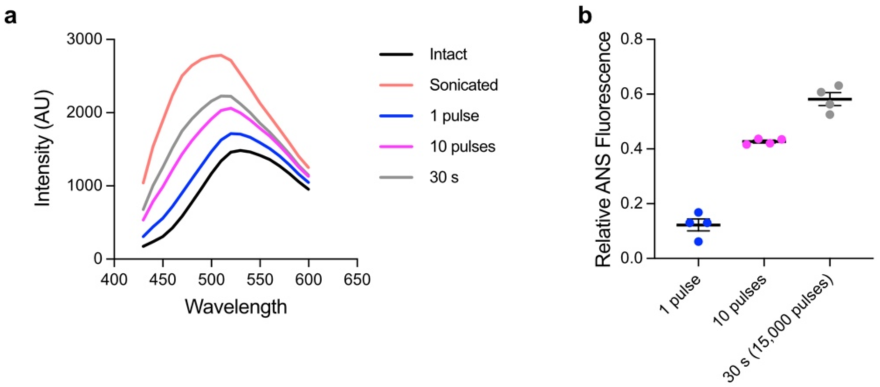


**Figure S9.** *Anabaena* GVs collapse under FUS insonation. **a)** Representative emission spectra of OD 0.25 *Anabaena* GVs with 100 µM ANS following insonation at 0.65 MPa PNP, 500 Hz pulse repetition frequency. **b)** Relative ANS fluorescence at 480 nm of GVs after exposure to 1, 10, or 15,000 FUS pulses at 0.65 MPa PNP, normalized to fluorescence of intact and pre-collapsed samples. *N* = 4. Error bars, ± SEM.

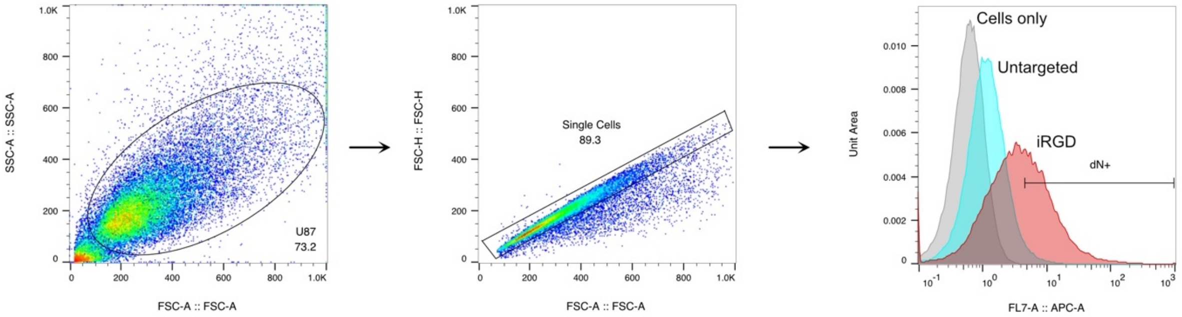


**Figure S10.** Flow cytometry gating strategy for iRGD targeting experiments. Debris was excluded based on SSC-A vs FSC-A. Single cells were selected from FSC-H vs FSC-A. Bicone positive cells (dN+) were defined relative to a cell-only control sample.

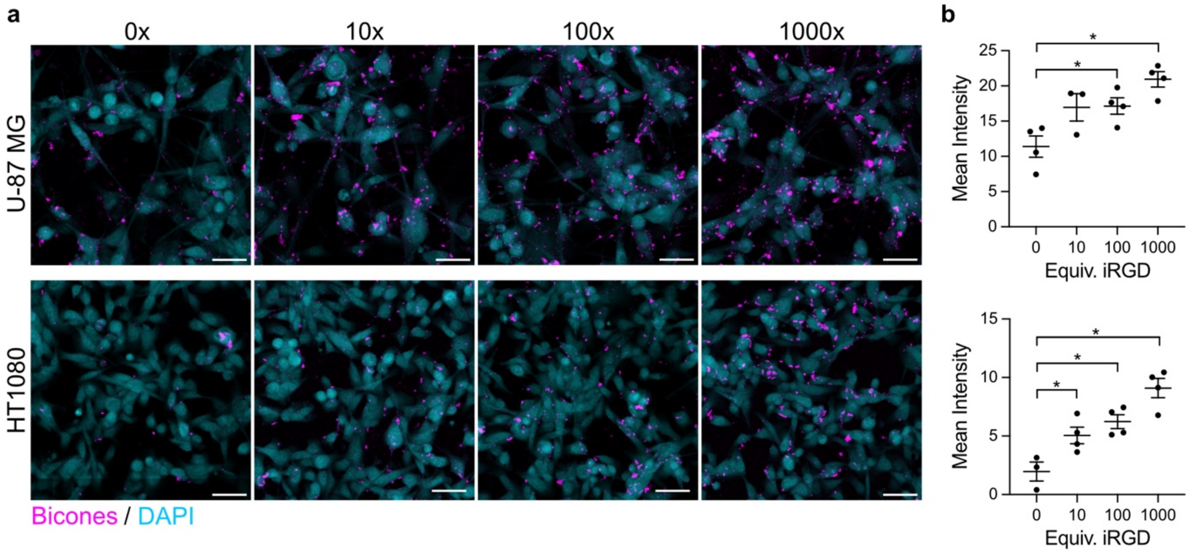


**Figure S11.** Targeting improves with iRGD loading. **a)** Representative confocal microscopy images of cells after incubation with bicones carrying the indicated maximum stoichiometry of iRGD peptides. Scale bars, 50 µm. **b)** Mean fluorescence intensity in the bicone channel of cell regions in images from panel A. U-87 MG (top) and HT-1080 (bottom) cells were defined by thresholding on autofluorescence in the DAPI channel. *N* = 4. Error bars, ± SEM. Welch's t-test, (*, $p < 0.05$).

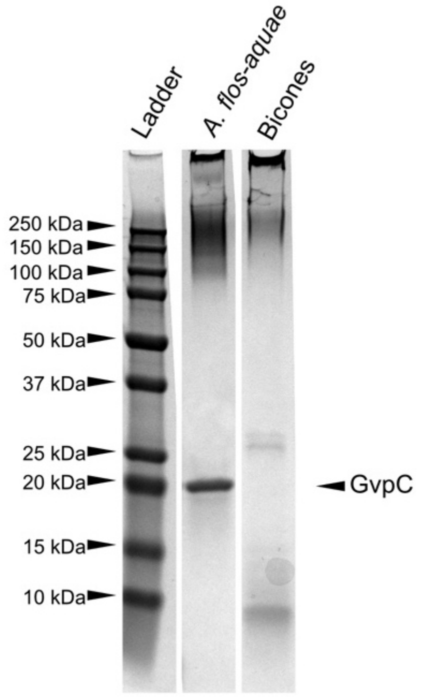


**Figure S12.** SDS-PAGE of bicones (200 µg mL$^{-1}$) and *Anabaena* GVs (OD 10). GvpC (indicated with an arrow) is not found on bicones.

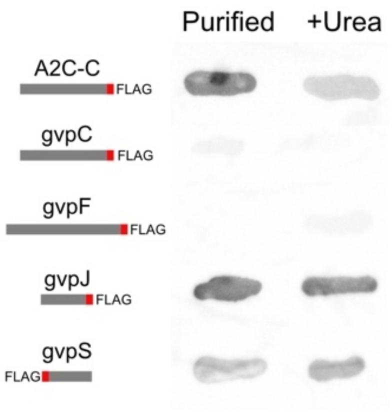


**Figure S13.** Anti-FLAG dot blot of purified bicones containing the indicated FLAG-fusion protein before and after treatment with 6 M urea to remove surface-bound proteins.

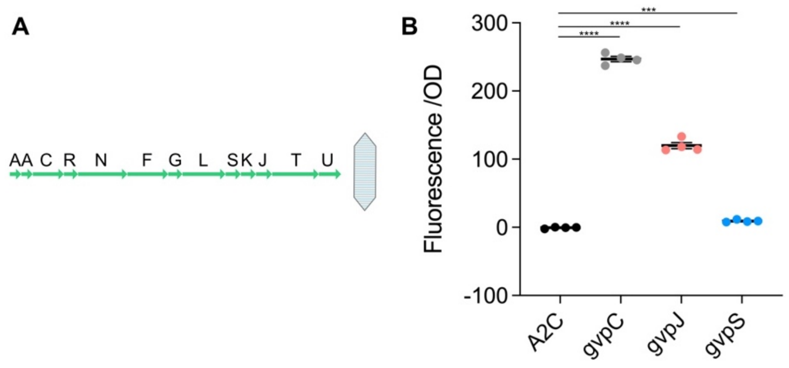


**Figure S14.** GvpJ and GvpS can also be used to genetically functionalize larger GVs. **a)** A2C gene cluster. The SpyTag peptide was appended to the end of gvpC, gvpJ, or gvpS. Purified particles were then reacted with SC-mNG. **b)** Fluorescence of purified samples after conjugation. *N* = 4. Error bars, ± SEM. Welch's t-test (***, p < 0.001; ****, p < 0.0001).

| | **Bicones** | **Anabaena** |
|---|---|---|
| Length (nm) | 72 | 519 |
| Diameter (nm) | 40 | 85 |
| Total volume (aL) | 0.03 | 2.784 |
| Gas vol/total vol | 0.764 | 0.903 |
| Particles/mg protein | 1.01e14 | 2.65e12 (~OD27 in 1 mL) |
| Acoustic Collapse Midpt (MPa) | ~2.4 | ~0.9 |
| Hydrostatic Collapse Midpt (MPa) | ~1.2 | ~0.6 |

**Table S1.** Physical properties of bicones compared to *Anabaena* GVs.

| | | Forward | Reverse |
|---|---|---|---|
| gvpC | SpyTag | TAAGCCGACGAAGGGAGGATCTGGGATTTCTTTAATGGCAAAAATCC | tatgcgtctaccattacaatatgtgcCATGAAGTTCTCCAAAAAATAG |
| | FLAG | gataaagggggctctgggATTTCTTTAATGGCAAAAATCC | gtcatcatctttataatcCATGAAGTTCTCCAAAAAATAG |
| | deletion | GAGAACTTCTAAAGATCTAACTATTGGAGGCTACTAAAAATG | CAAAAAATAGTAAATTAGCGCTAGCAAG |
| gvpR | SpyTag | | |
| | FLAG | gataaagggggctctgggGAAATTAAAAAAATTATGCAAGC | gtcatcatctttataatcCATTTTTAGTAGCCTCCAATAG |
| | deletion | GCGATAAGATGGCAGGAGCTTG | CTTTAGTAGCCTCCAATAGTTAGATCTTTATTAACC |
| gvpF | FLAG | gatgatgacaaaTAACGTGCTTCACAAATTAG | gtctttataatccccagagcccccTTTCTCTTCTACTTTTAGGC |
| | deletion | cgtgcttcacaaattagtaaccgc | GTTTTCAAGCTCCTGCCATCTTATC |
| gvpG | FLAG | aaagacgatgatgacaaaTAGATGGGAGAATTACTG | ataatccccagagcccccGGATTCCTCATTTCTTTTTTG |
| | deletion | GAGAAATAAATGGGAGAATTACTGTATTTATACGG | CTCTACTTTTAGGCGAATGTTCAC |
| gvpL | FLAG | gataaagggggctctgggGGAGAATTACTGTATTTATACG | gtcatcatctttataatcCATCTAGGATTCCTCATTTC |
| | deletion | CGTGAGGAATTAACATTATGTCTCTTAAAC | CTAGGATTCCTCATTTCTTTTTTGTGTTAGCTCTTC |
| gvpS | SpyTag | TAAGCCGACGAAGGGAGGATCTGGGTCTCTTAAACAATCCATGG | tatgcgtctaccattacaatatgtgcCATAATGTTAATTCCTCACTTTAC |
| | FLAG | gataaagggggctctgggTCTCTTAAACAATCCATGG | gtcatcatctttataatcCATAATGTTAATTCCTCACTTTAC |
| | deletion | gatgcaaccggtcagc | gtgttaattcctcactttacgc |
| gvpK | FLAG | GATGATGACGATAAATAAGCGGTCAGTAGGAGGAAC | CTTGTAATCCCCAGAGCCCCCAAGCAGGCTGCCTAGCGG |
| | deletion | cggtcagtaggaggaacag | gtcaggatccaagtggattcg |
| gvpJ | SpyTag | GTAGACGCATATAAGCCGACGAAGTAAAAACTGTACGCTACTTAAAAAA[illegible] | CATTACAATATGTGCCCCAGATCCTCCACGTTTCGTTTCTATTTTT[illegible] |
| | FLAG | aaagacgatgatgacaaaTAAAAACTGTACGCTACTTAAAAAATG | ataatccccagagcccccACGTTTCGTTTCTATTTTTTC |
| | deletion | gaactgtacgctacttaaaaaatg | cctgttcctcctactgac |
| gvpT | FLAG | gataaagggggctctgggGCAACTGAAACAAAATTAGATAAC | gtcatcatctttataatcCATTGTAAATCCCTCCATTTTTTAAG |
| | deletion | gacgtaaaggaggaaagaaag | ggtaaatccctccatttttaagtag |
| gvpU | FLAG | gataaagggggctctgggAGTACAGGCCCTTCTTTTTC | gtcatcatctttataatcCATGTCTTTCTTTCCTCCTTTAC |
| | deletion | GTCTTTCTTTCCTCCTTTACGTC | CAAACGGCGGGGTGATTGC |

**Table S2.** Primers used for appending SpyTag and FLAG tags, or for gene deletion.
